# Neural responses to natural visual motion are spatially selective across the visual field, with selectivity differing across brain areas and task

**DOI:** 10.1101/2021.03.05.434148

**Authors:** Jason J Ki, Jacek P Dmochowski, Jonathan Touryan, Lucas C Parra

## Abstract

It is well established that neural responses to visual stimuli are enhanced at select locations in the visual field. While spatial selectivity and the effects of spatial attention are well-understood for discrete tasks (e.g., visual cueing paradigms), little is known about neural response during a naturalistic visual experience that involves complex dynamic visual stimuli, for instance, driving. In this study, we assess the strength of neural responses across the visual space during a kart race video game. Specifically, we measure the correlation strength of scalp evoked potentials with optical flow magnitude at individual locations on the screen. We find the strongest neural responses for task-relevant locations in visual space, selectively extending to areas beyond the focus of overt attention: while the driver’s gaze is directed upon the heading direction at the center of the screen, we observe robust neural evoked responses also to peripheral areas such as the road and surrounding buildings. Importantly, this spatial selectivity of neural responses differs across scalp locations. Moreover, during active gameplay, the strength of the spatially-selective neural responses are enhanced compared to passive viewing. Spatially selective neural gains have previously been interpreted as an attentional gain mechanism. In this view, the present data suggest that different brain areas focus attention on different task-relevant portions of the visual field, reaching beyond the focus of overt attention.

## Introduction

Traditional studies of visual perception employ tightly controlled experimental paradigms. For instance, classic behavioral studies on visual attention present discrete stimuli and attention cues at select areas in the visual field and measure accuracy or response times as a function of location and cues (Posner, 1980). These studies have established a clear difference between overt attention, defined as the location of observable gaze position, and covert attention, which manifests as a performance gain when subjects are given a cue directing their attention to a location different from their gaze position (Moran and Desimone, 1985; Spitzer et al., 1988). Studies on the effects of covert attention on neural response often present discrete stimuli at specific locations in the visual field and evaluate the effect of attentional cues on neural activity (McAdams and Maunsell, 1999; Motter, 1993). Using intracranial recordings, animal and human studies have established that neuronal firing to visual stimuli is selectively enhanced for attended locations in the visual field (Luck et al., 1997; Moore, 1999; Self et al., 2016). Similarly, location-dependent neuronal gains have been found for attended locations with scalp recordings in humans. For instance, discrete visual stimuli produce robust contralateral responses when covertly attending to a selected visual hemisphere (Hillyard and Anllo-Vento, 1998; Luck et al., 1990; Mangun, 1995). Moreover, similar results have been shown in neuroimaging studies with the enhanced contralateral hemodynamic response over the visual cortex during spatial attention tasks (Beauchamp et al., 2001; Mangun et al., 1998; Tootell et al., 1998).

While a great deal has been learned from these studies, real-world tasks such as driving differ considerably from these traditional experimental paradigms. Starting with posner visual cueing paradigms (Posner, 1980), during studies of covert attention (e.g. (Luck et al., 1990; Mangun, 1995; Moran and Desimone, 1985; Rugg et al., 1987; Spitzer et al., 1988), subjects are often asked to inhibit saccades. Given the unnatural task constraints, it is unclear how attention is deployed in visual space during ecologically valid tasks. Existing evidence from evoked potentials already suggests that neural processing is altered when saccades are artificially constrained (Ki et al., 2016; Kulke et al., 2016). Moreover, traditional paradigms employ isolated stimuli, which are often presented at discrete locations in space or discrete moments in time (Hillyard and Anllo-Vento, 1998; Mangun et al., 1998). In contrast, day-to-day visual experiences involve dynamic motion with visual inputs that are spatially-heterogeneous. Thus, given the complexity of naturalistic stimulus, alternative experimental paradigms and analysis methods are necessary, if only to validate theories of attention developed under the reductionist approach.

In recent years, data-driven methods have developed to quantify the neural activity generated in response to complex naturalistic stimuli. These studies employ multivariate modeling to map low-level stimulus properties to observed neural response (encoding) or vice-a-versa (decoding) (Holdgraf et al., 2017; Naselaris et al., 2011). This stimulus-response modeling is also referred to as system identification ((Gallant et al., 2012; Wu et al., 2006), and is particularly advantageous for complex dynamic visual stimuli, which can not be readily decomposed into discrete events required for conventional event-related analysis (Crosse et al., 2016). One area of research that has popularized the stimulus-response modeling approach is studies on visual representation using fMRI. These studies capture spatial, and orientation features of naturalistic images and dynamics movies to predict hemodynamic response at individual voxel regions (Kay et al., 2008; Naselaris et al., 2009; Nishimoto et al., 2011), and the mapping can be reversed to reconstruct the original stimulus with reasonable accuracy (Miyawaki et al., 2008; Naselaris et al., 2009; Nishimoto et al., 2011). A basic finding is that; different regions of the visual cortex exhibit spatial selectivity to visual input at different locations across the visual field, with selective attention increasing the reliability of these responses (Kay and Yeatman, 2017).

In electrophysiology, similar encoding models have been applied to map temporal visual contrast to neural responses (Gonçalves et al., 2014; Lalor et al., 2006; Lalor and Foxe, 2009). This approach yields a precise estimation of the evoked response dynamics, which is analogous to event-related potentials; however, it does not account for the spatial heterogeneity of visual dynamics. In a similar approach, the strength of evoked activity across multiple electrodes can be measured as the correlation of stimulus and response (Dmochowski et al., 2018). Using this approach we previously investigated the effects of active perception by comparing the strength of neural response during active play and passive viewing of a videogame (Ki et al., 2020). We found that active engagement in the task enhanced responses to the overall visual contrast dynamic.

Based on the spatial selectivity and attention effects observed in EEG and fMRI, we hypothesized that during a natural visual task, the strength of neural responses differs across the visual field and that the task selectively modulates neural activity. To test this, we re-analyze the EEG recordings from (Ki et al., 2020) in which subjects either actively played a kart racing video game or passively viewed a pre-recorded video of a race. We extended the system identification approach by measuring stimulus-response correlation resolved in visual space. In doing so, we model response to a natural dynamic visual stimulus while capturing spatial selectivity not typically observed with EEG. Our findings show that the neural response to movement is enhanced in task-relevant areas of the visual scene. Remarkably, this spatial selectivity differs across brain areas. Moreover, we find that active play enhances neural responses in areas extending beyond the focus of overt gaze position. Overall, these results suggest that neural response during a naturalistic dynamic visual experience differs across the visual field depending on task demands.

## Results

Human subjects (N=42) were asked to play a kart race video game (Fig. 1), while we recorded continuous EEG activity and in some subjects also gaze position (N=17). The players control velocity and left/right heading direction with their right hand. Their goal is to complete the course as fast as possible (each race approximately 3 minutes to complete). Good performance implies driving fast without veering off the road or crashing into buildings and competing race karts, which incurs a large time penalty. The visual dynamic is dominated by the translation of the moving vehicle, which produces an outward radial pattern of optical flow (Fig. 1A), computed here for each screen location (Horn and Schunck, 1981). The optical flow across all races is dominant in the periphery and weakest at the center (Fig. 1B). In contrast, the task-relevant information is on the road and adjacent buildings (see [video clip](https://drive.google.com/file/d/1xt7OvM2xuVep_C-MEGd9cmqnQ3XHMxvO/view), eye position markers: cyan circle - active play, green circle - passive viewing). Interestingly, judging by the distribution of gaze positions (Fig 1C), overt attention seems to be narrowly focused on the road ahead. During the kart race, the gaze distribution is similar to that of real-world vehicle control in which the driver’s eye gaze is particularly focused on the inner ‘tangent point’ of the road curvature [Land & Lee, 1994]. The right-ward skew of the eye gaze distribution (Fig. 1C) indicates that gaze is directed at this curvature point.

**Figure 1.**
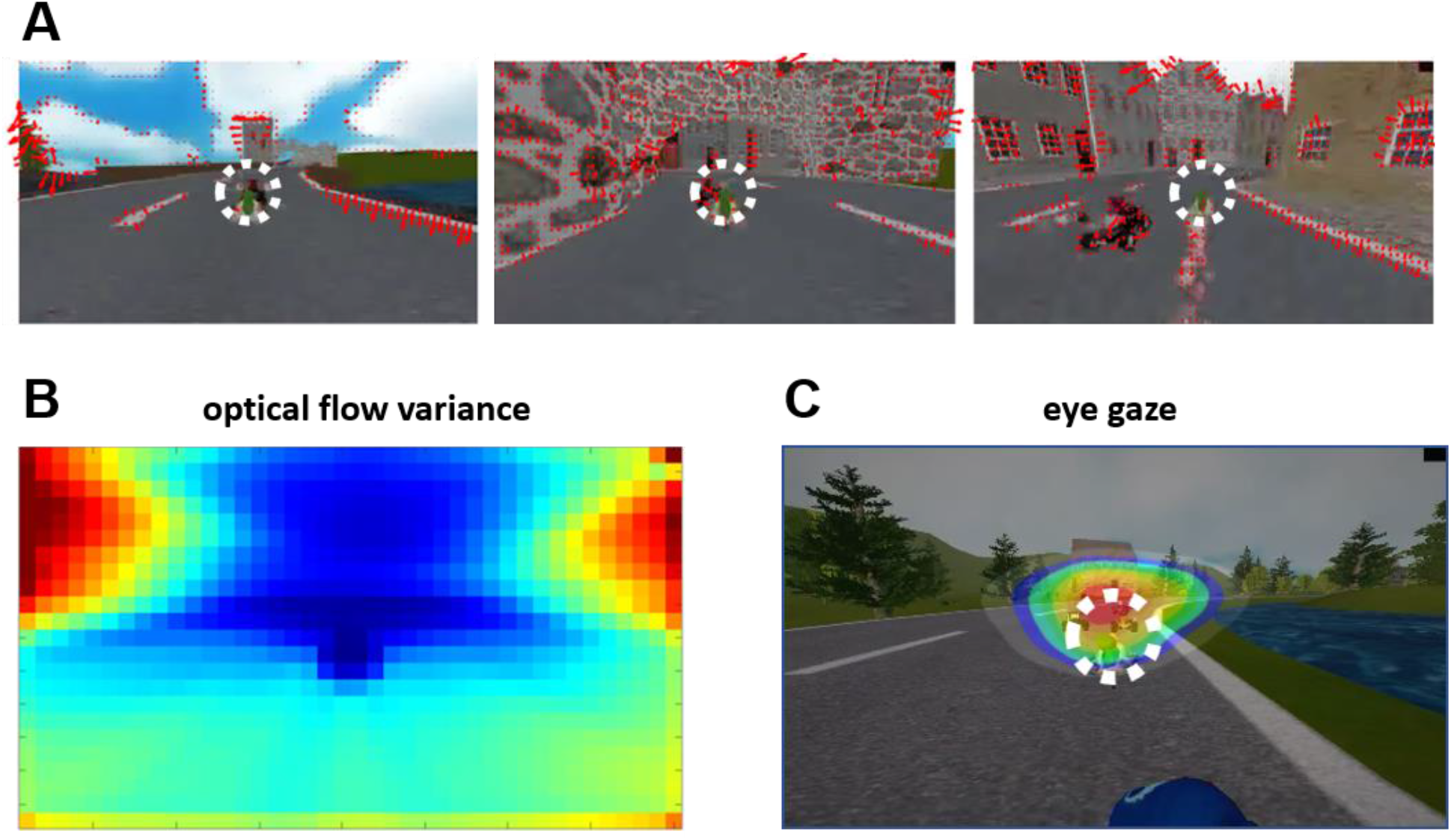
Visual dynamics of the race kart video game. A) The kart’s translation creates the overall visual dynamics of the racecourse, which forms optical flow that expands radially outward. B) On average, visual flow is slow in the center and faster at the edge of the screen, which is demonstrated by the high variance of optical flow in the lateral region. C) Subjects’ eye gaze distribution is at the center of the screen near the road’s curvature, mainly focused at the tangential point towards the general turning direction (the right band of the contour plot indicates the counterclockwise navigation around the track).

### Spatially-Resolved Stimulus-Response Correlation

Given that the player’s kart is centered on the screen (Fig. 1A, dotted circle), absolute locations on the screen take on distinct importance in this driving task. We hypothesized that the strength of the neural response to visual movement is enhanced at select areas on the screen that are task-relevant. To test this, we assess the strength of visually evoked activity resolved in visual space. Specifically, we measure stimulus-response correlation (SRC) between the raw EEG signal and optical flow magnitude at individual screen locations (Fig. 2). We will use two different approaches to measure SRC. In the first approach, the multiple EEG electrodes are projected onto a component space (Dmochowski et al., 2018) similar to the principal or independent components -- these can be conceived as “virtual electrodes” -- and the stimulus is filtered to predict the neural activity observed in these virtual electrodes (Fig. 3). In the second approach, the stimulus is filtered to directly predict neural activity in each of the original EEG electrodes [Lalor 2006] (Fig. 4). In both instances, the temporal filters -- i.e., impulse responses or temporal response functions (TRF) -- are estimated for each screen location separately, and SRC is the correlation of the predicted neural activity with the actual neural activity. Given that optical flow for this video game is sufficiently distinct at each of the screen locations (Fig. 2A), one might expect quite different SRC for each location (Fig. 2C).

**Figure 2.**
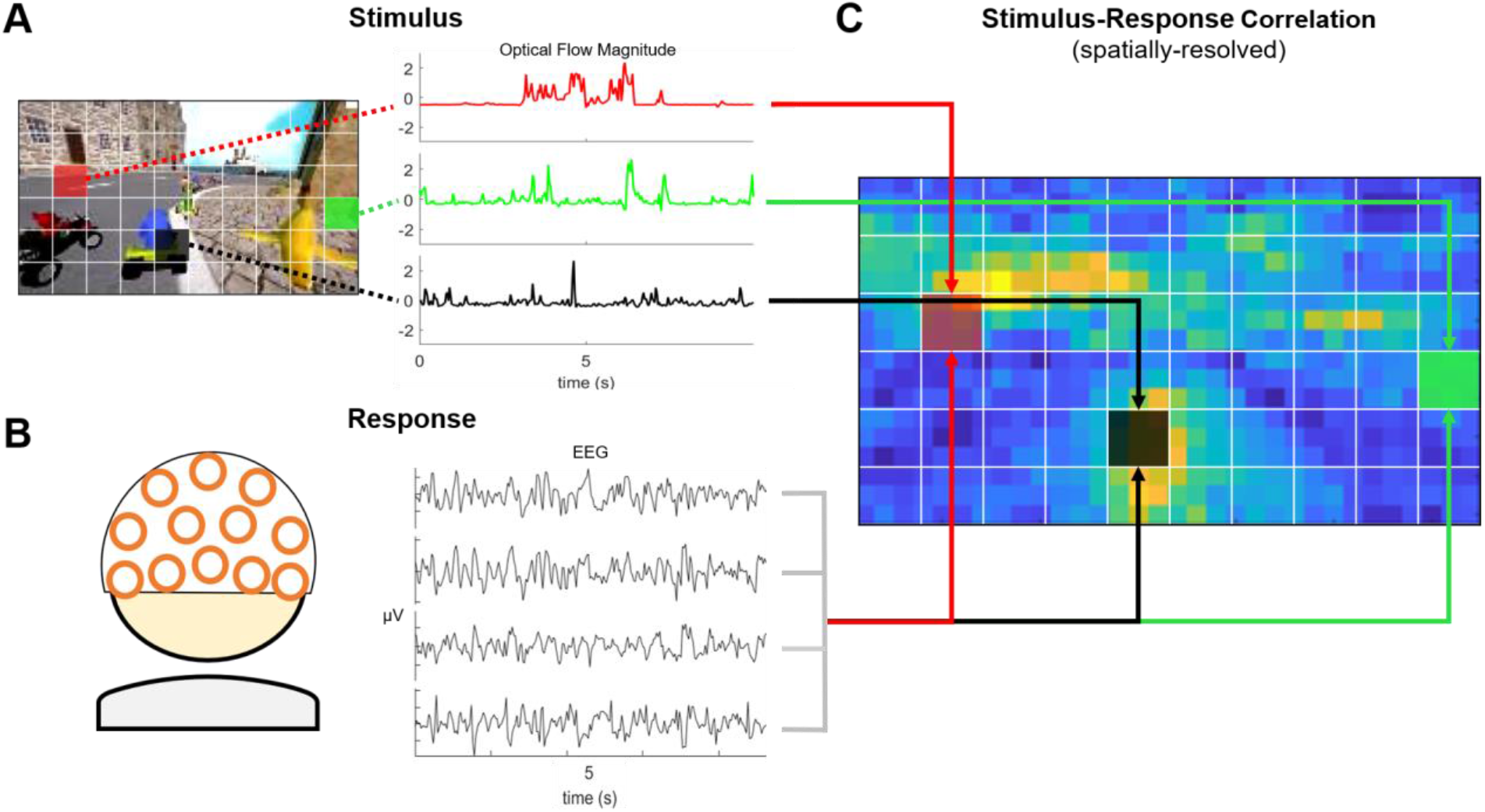
Resolving the strength of neural response across visual space. Here, we illustrate the spatially-resolved stimulus-response correlation (SRC) approach, which measures the correlation of continuous-stream neural response to a temporally filtered stimulus. **A)** We record the screen capture of the video game and extract the optical flow at each location on the screen (Horn-Shunck 1981). The image shows a snapshot of optic-flow magnitude. Optical flow is averaged over small image patches (7×8.5 pixels). The colored patches at different screen locations (red, green, black) show unique optical flow magnitude over time. **B)** Neural responses are captured with the scalp EEG during the videogame presentation. SRC is computed as the correlation of optical flow magnitude with the neural response, for which we use either individual electrodes of EEG or virtual electrodes (i.e., spatial projection of multi-channel EEG). **C)** We assess the location-dependence of neural response by measuring the correlation between the EEG and temporally filtered optical flow magnitude at individual patch regions (23×36, screen capture recorded 180×320 pixels).

**Figure 3.**
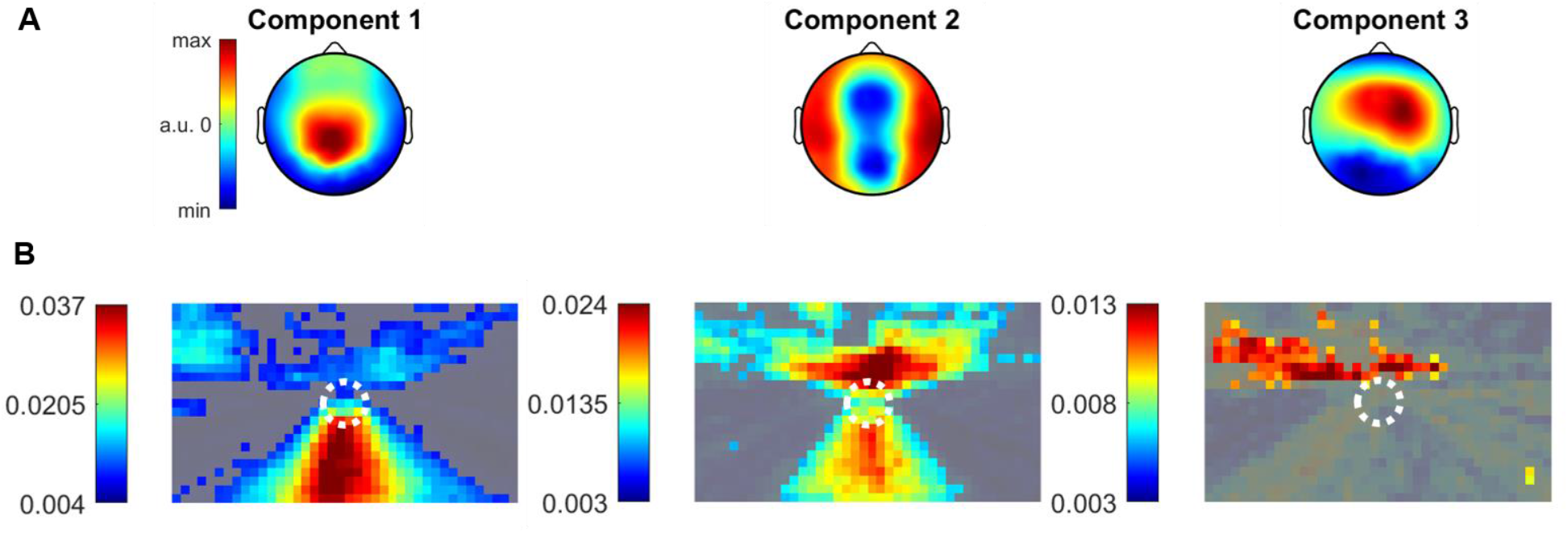
The strength of the visually evoked response is location-dependent. **A)** For the spatially-resolved analysis, we employ the spatial and temporal filters previously computed in Ki et al. 2020 using canonical correlation analysis (CCA). CCA yields a set of components that maximize the correlation between spatially filtered EEG (96 channel) and temporally filtered stimulus. Components of the EEG can be thought of as virtual electrodes, and the associated stimulus filters correspond to temporal response functions (shown in Fig. S1). **B)** We measure the correlation between spatially filtered EEG and temporally filtered optical flow at individual screen locations (24 x 36 patches) for the top 3 CCA components. Here, we create a visualization of spatially-resolved SRC using a colormap. The color value at each pixel maps to SRC based on a color scale. The areas with grey show patch regions in which SRC is below chance correlation (p > .05).

**Figure 4.**
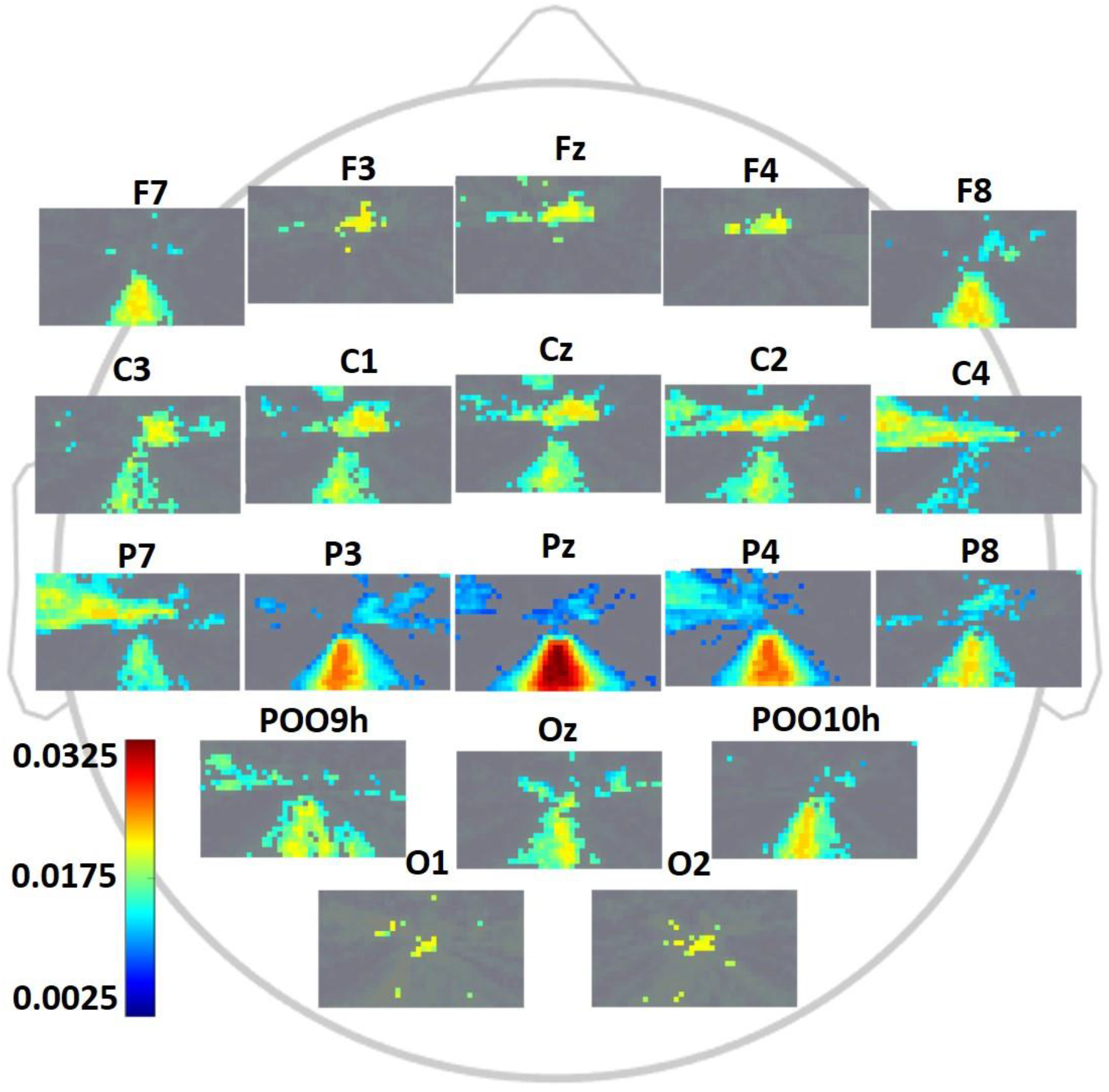
Location-dependent neural responses differ across scalp positions. Stimulus-response correlation computed from optic flow at each screen location with each electrode. SRC indicates the correlation between actual EEG and EEG predicted by filtering optic flow magnitude with a temporal-response function. The areas with grey show patch regions in which SRC is below chance correlation (p > .05).

### The strength of the visually evoked response varies across visual space

We analyzed the strength of stimulus-response correlation (SRC) for different locations across the screen (23 × 36 patches) for all subjects and races combined (each subject performed two races in active play or passive viewing conditions, yielding a total of n = 170 races, 179.91s +/− 17.14s duration each). Temporal correlations are measured at the 30Hz frame rate of the video (EEG is downsampled to this rate).

To quantify the overall SRC, we first used components of the EEG that were optimized to capture the strongest correlation between global optic flow (mean amplitude over the entire screen) and EEG responses following (Ki et al., 2020). Specifically, n=96 EEG electrodes are combined linearly, generating an EEG component signal which can be thought of as virtual electrodes (see Methods). The corresponding distribution of these components (Fig. 3A), is known as “forward model” (Haufe et al., 2014). Temporal response functions (TRFs) to predict the EEG component activity are then estimated for each patch on the screen separately (Fig. S1). The dependence of these TRFs on visual field and scalp locations will be discussed later (in Fig. 6). Stimulus-response correlation (SRC) is the correlation between the observed EEG, in each component, with the activity predicted from optic flow in each screen location (Fig. 3B). We consider the top three components as they capture most of the SRC (Notice in the colormap that SRC diminishes for the later components). Stronger SRC indicates stronger neural response to local optic flow. For component 1, which captures the strongest neural response, SRC is highest on the road. The optic flow in this area is dominated by road demarcation as well as the movement of competing cars and other objects. In component 2, the neural response is strongest for movement on the horizon where buildings appear as well as the road. In component 3 (right-lateralized scalp topography), the neural response focuses on the horizon, primarily to the left visual hemisphere. These unique spatial patterns suggest that the strength of visual evoked responses varies with location on the visual field, but also, this spatial preference differs for different brain areas.

### Visual-spatial dependence of neural responses varies across EEG electrodes

To determine specifically how the visuo-spatial selectivity of SRC differs across brain areas, in the second approach, we used TRF to map the stimulus to the neural response directly for each scalp electrode. The TRF estimation follows an established system identification approach (Crosse et al., 2016; Lalor et al., 2006), except that we again resolve SRC in visual space by computing unique TRF from the optic flow at individual screen locations (Fig. 4, showing a subset of 20 of the 96 electrodes). For midline-frontal channels, SRC is strongest at the top center of the screen, yet, lateral-frontal electrodes respond strongest to movement on the road. At central electrodes, over cortical somatosensory and motor areas, we observe responses lateralized to the contralateral visual field (which is dominated by the movement of buildings that are to be avoided). Note that responses are stronger in the left visual hemifield, but this dominance largely disappears in the passive viewing condition (Fig. S2), suggesting that this is the result of the right-hand key presses to control the kart. At parietal electrodes, we observe strong responses to visual dynamics on the road. This parallels the finding for component 1 (Fig. 3), which had a dominant parietal positivity. Lastly, in the occipital electrodes, we observe a strong response to visual dynamics associated with movements on the road, likely associated with competing karts in the vicinity of the player’s vehicle or objects on the road. Finally, overall there is a predominant bias to the left hemifield, which is also expressed in component 3. Overall, these results show that spatial selectivity of neural response is different across different brain areas.

### Neural response is enhanced at task-relevant locations during active play

The manual control of the race kart involves active engagement, requiring selective visual-spatial attention to task-relevant areas. Given the modulation of neuronal gain observed with visual attention (Hillyard et al., 1998; Luck et al., 1997; Maunsell and Cook, 2002; Motter, 1993), we hypothesize that the EEG response will be selectively enhanced during active play for locations that are relevant to the task. To test this, we compared the strength of neural responses to optic flow between active play with passive viewing. In the passive viewing condition, subjects were asked to watch prerecorded races of the game. The order of active play (n = 84 trials; a trial is three laps around the racecourse) and passive viewing (n = 86 trials) was counterbalanced across subjects (N = 42). We compute the location-dependent SRC as in Figure 3. To obtain an assessment of the overall EEG response, we summed SRC over the top 3 components (Fig. 5). In Figure S2, we resolve this by individual electrodes. The mean across subjects has distinct spatial distribution across the scene for active and passive play (Fig. 5A). During active play, neural responses are more robust for movement occurring on the road and beyond the road where buildings and trees could result in collisions. In contrast, during passive viewing, the neural response is strongest only for areas covering the road. Indeed, we find that neural response is significantly enhanced with active play for areas that cover buildings and objects coming ahead on the horizon above the road (Fig. 5B highlights locations with p > .05, Wilcoxon rank-sum, one-tailed, FDR corrected, N=42). This enhancement extends beyond the focus of overt attention as characterized by gaze position (Fig. 5C). Notably, the contrast seems to be driven mostly by central and frontal scalp locations (Fig. S2). Importantly, we did not find differences in gaze position between active play and passive viewing (Fig. 5D highlights locations with p > .05, Wilcoxon rank-sum, one-tailed, FDR corrected, eye-tracking data was available in N = 17 subjects). The distribution of eye velocity appears to be slightly skewed towards larger saccades in active play (Fig. 5E). However, pair-wise comparison of the average velocity of individual subjects between active and passive trials showed no significant difference (Fig. 6F, z = 0.355, p = 0.72, df = 16, paired two-tailed Wilcoxon signed-rank test; i.e. only including subjects for whom both conditions were available) indicating that eye movement dynamics are comparable in the two conditions. Overall, these results suggest that the neural response is enhanced at task-relevant locations, extending beyond the focus of overt attention, and differences cannot be attributed to changes in fixation position of saccade size.

**Figure 5:**
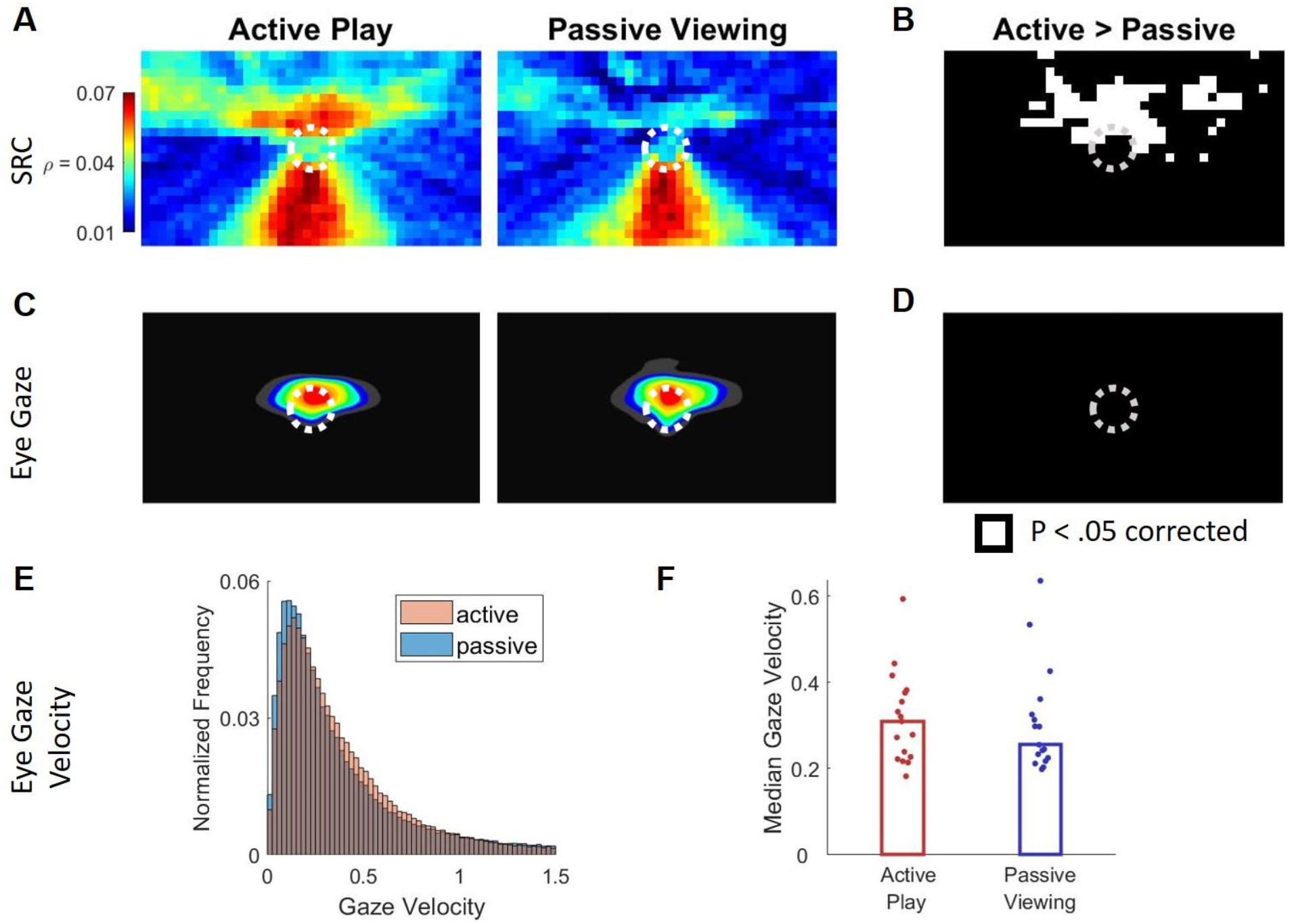
Neural response to optic flow is enhanced at task-relevant locations during active play, while overt gaze position and saccade velocity are unchanged: **(A)** Spatially resolved SRC (as in Fig. 3) summed over three components but separate for active play and passive viewing conditions. **(B)** Locations of significant differences in SRC between active and passive conditions. Enhancement occurs are task-relevant locations. **(C)** Distribution of gaze position over time-averaged over subjects. The gaze is mostly focused on the heading direction above the stationary position of the subject’s own kart (white dotted circle). **(D)** The contrast in the distribution of gaze position between active play and passive viewing shows no statistical differences (P > .05). **(E)** Distribution of gaze velocity suggests a slight increase in gaze dynamics during active play. **(F)** Median velocity of individual subjects, however, shows no significant difference between conditions (each point is a subject).

**Figure 6.**
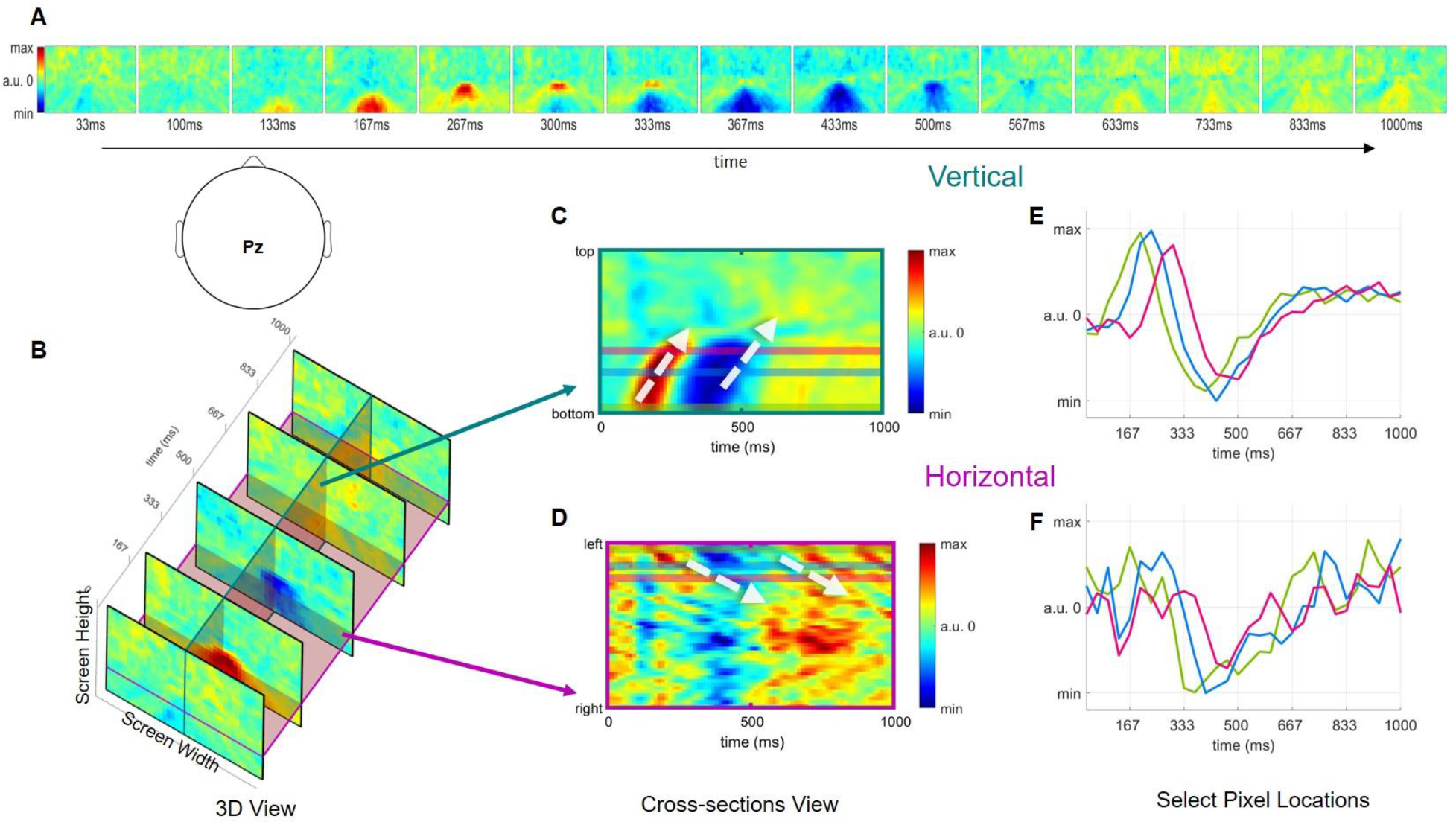
Neural response to peripheral movement is faster. (**A**) Spatially-resolved temporal response function (TRF) at select time points. The TRFs are computed by regressing optical flow at each screen location with Pz electrode. **C**) 3D visualization of spatially resolved TRFs. The slice along the time dimension indicates cross-sections of temporal response at vertical (Dark Green) and horizontal (Maroon) positions on the screen. (**C**) TRF along vertical screen direction. White dashed arrows indicate increasing delay towards the center of the screen (opposite to the direction of optic flow, which moves from the center outwards) (**D**) TRF in horizontal screen direction. (**E**) Time course at select pixel positions (green, blue, pink), which corresponds with colored lines along the vertical cross-section in C. (**F**) Same as panel E but along the colored lines at the horizontal cross-section.

### Neural response occurs faster at the periphery

To determine if the latency of neural responses differs across the visual space analyze the TRFs that were extracted separately for each visual location and electrode (for Fig. 4). Analogous to ERPs, TRF yields an estimate of evoked-response time to a specific visual input. Here, we focused on the parietal Pz electrode and displayed the distinct TRFs obtained by regressing optical flow at each 2D screen location (Fig. 6A). In combination, the collections of TRFs have two visual space dimensions and one time dimension indicating the latency of neural response to optical flow at each screen location (Fig. 6B). We find a dependence of latency on location in visual space, which is most obvious when displaying the TRFs in time along a vertical direction (Fig. 5C, indicated by the white dashed arrows). Neural response first peaks at 130ms for movement at the bottom of the screen, with latency increasing towards the center of the screen, where it reaches a latency of 300ms (Fig. 6E, arrow). A similar pattern is observed for electrodes Cz and Oz (Fig. S3) as well component 1 (Fig. S1). This delay is primarily driven by the movement on the road. In the horizontal direction extending beyond the road, we also see a similar phenomenon albeit with different latency and less pronounced (Fig. 6D). Notably, the delay does not seem to depend on whether the subject is actively engaged in the game (Fig. S4). In summary, in both horizontal and vertical cross-sections, the peak of neural response occurs first at the lateral positions of the screen and finishes last at the center, and this effect is not dependent on task engagement.

## Discussion

In this study, we assessed the strength of neural responses elicited by stimuli across the visual space during a dynamic video game. The task involved racing a kart while avoiding obstacles, as collisions significantly impact race time, which was the primary objective of the game. This task requires selective attention to specific objects or locations on the screen in order to plan and control the kart. Because the kart location was maintained at the center of the screen, and obstacles appeared in similar locations on the screen, we can draw inferences on the task-relevance of neural responses to movement occurring at each location of the screen. The need for spatial consistency is perhaps a caveat of the approach; nonetheless, it allows us to derive a visual-spatial representation for EEG, which has traditionally been constrained to the temporal domain.

Similar to real-world driving (Land and Lee, 1994), we found that subjects’ gaze was fixated on the curvature of the road while simultaneously shifting the gaze to competing karts and objects. The translation of the kart along the road produces a uniform optical flow that radiates outwardly, while other competing karts produce sparse optical flow along the road. Due to this self-motion, the optic flow was strongest in the lateral periphery (Warren and Hannon, 1988). Yet, we find that neural responses were strongest for movement in the middle of the screen, including the road and obstructions coming up on the horizon. Therefore, neural responses are strongest for task-relevant locations. Importantly, this spatially selective neural response extended beyond the focus of gaze position, which is the established marker of overt visual attention.

The spatial selectivity differed across scalp locations. For example, parietal electrodes show strong responses for the road, which is dominated by the movement of traffic lines demarcating the road and competing karts. It is worth noting that visuomotor transformations, as may be required in this task, are known to occur in the superior parietal cortex, as demonstrated in non-human primates (Fogassi and Luppino, 2005) as well as parietal components of human EEG recordings (Naranjo et al., 2007). The parietal activity responded earlier to movement in the periphery and later to movement at the center of the screen. This is intriguing as it is in the opposite direction of objects moving from the center to the periphery. The effect was most pronounced along the road. Earlier studies on the latency of evoked potentials as a function of visual eccentricity in static visual stimuli give mixed results with latency depending on upper vs. lower and contralateral vs. ipsilateral hemifield (Busch et al., 2004; Capilla et al., 2016). During natural vision, different regions are selective to local motion and global flow (Bartels et al., 2008). Given that the perception of self-motion is mainly determined by the magnitude of visual flow in the periphery (Warren et al., 1988), it is possible that peripheral visual motion is processed faster in the brain. Processing of optic flow is ascribed to the dorsal visual stream, particularly in the Medial Superior Temporal (MST) areas (Duffy, 1998; Tanaka and Saito, 1989) which may have contributed to this component. An alternative interpretation, given the relatively long response onset time of 200-300ms, is that visual content in the center of the screen had more complex semantics and therefore resulted in longer processing times, consistent with the general finding that visual processing to high-level properties of the stimulus (VanRullen and Thorpe, 2001). For instance, ERP studies show that spatial attention is deployed faster than attention to visual features (Anllo-Vento and Hillyard, 1996; Liu et al., 2007).

Neural responses were significantly enhanced during active play of the game compared to passive viewing of the same visual stimuli. This enhancement was spatially selective, overlapping with the focus of overt attention at the center of the screen, but also extending to task-relevant locations beyond. It is well-established that visual evoked potentials are enhanced when attending to a specific target or locations, even in the absence of overt orienting of eye gaze (Hillyard and Anllo-Vento, 1998; Luck et al., 2000; Luck and Kappenman, 2011; Mangun and Hillyard, 1988). Here similarly, neural responses were enhanced without a change in gaze position, suggesting that this may be the result of task-relevant increase of spatially selective attention.

We note that this selective spatial enhancement for active play was strongest for central electrodes over the motor cortex, which responded more strongly to visual movement on the horizon in the contralateral visual hemisphere (where upcoming buildings appear alongside the road). This left/right symmetry can not be ascribed to the neural activity associated with somatosensory or hand motor control, as driving the kart involved a single hand (the right hand). It is well-established that attending to one hemisphere of the visual field elicits stronger visual evoked potentials at the contralateral occipital cortex (Luck et al., 1990; Mangun, 1995). The visual-spatial selectivity of central scalp electrodes suggests that central brain areas also “attend” to visual movement on the contralateral visual field.

Another possibility is that the central components reflect an visuomotor integration required for navigating the kart. Note in this context that there was a dominance of responses to the left visual hemifield, notably stronger for the right hemisphere of the brain. This asymmetry largely disappeared during passive play. If this asymmetry was the result of somatosensory or motor evoked responses due to the actual right-hand key presses, one would have expected stronger asymmetry for the left (contralateral) brain hemisphere. This observation additionally supports the interpretation of motor areas “attending” to visual movement on the contralateral hemifield, something that may not have been reported previously in the literature. Note that even in the passive condition, there is a slight asymmetry. This may be the result of a slight asymmetry in the visual stimulus itself resulting from the predominant rightward curvature of the racetrack loop, which results in somewhat stronger optic flow on the left visual periphery, as well as the slight rightward skew of the gaze distribution as subjects focus on the upcoming road.

An alternative interpretation is that enhancement with active play results from bottom-up stimulus-driven reorienting, consistent with the observation that visually salient events shift eye gaze and activate dorsal ventral regions (Nardo et al., 2011). Indeed, a caveat of the present study is that during free viewing, eye position and eye movements are dynamic and uncontrolled. Traditionally, a distinction is made between overt and covert attention in experiments that aim to strictly control eye movements (Kelly et al., 2010; Kulke et al., 2016; Posner et al., 1984; Rugg et al., 1987). Eye movements such as saccades, microsaccades, pursuit, stigmas are distinguished as they contribute uniquely to attentional related ERPs (Kelly et al., 2010; Nobre et al., 2000). While we found that the overall eye movement velocity was quite similar between active and passive conditions, we did not analyze the effects of different types of eye movements in this investigation. It is possible that changes in eye movement contributed to enhanced neural responses. However, it has been shown that the magnitude of responses to visual cues remains similar regardless of eye movement condition (Parker et al., 2020). Moreover, covert and overt attention shifts elicit similar early evoked responses (Hanning et al., 2019). Taken together, we believe that the effects of differing eye movement dynamics played only a minor role, although we can not fully rule this out.

Methodologically, this study combines new and old approaches to studying neural responses to visual stimuli. Visual-spatial attention and feature-based attention have been extensively studied in the last 50 years using traditional event-related paradigms (Hillyard and Anllo-Vento, 1998; Luck et al., 2000; Luck and Kappenman, 2011; Mangun and Hillyard, 1988; Schoenfeld et al., 2007; Van Voorhis and Hillyard, 1977). More recently, decoding methods have been applied for continuous dynamic stimuli such as continuous speech, movies, and video games. (de Cheveigné et al., 2018; Dmochowski et al., 2018; Golumbic et al., 2013; Iotzov and Parra, 2019; Lalor and Foxe, 2010; Mesgarani et al., 2014; Mesgarani and Chang, 2012; Naselaris et al., 2011, 2011; O’Sullivan et al., 2015, 2015). The contribution of this study was to resolve neural responses in visual space while leveraging methods suitable to continuous dynamic stimuli. Thus, we believe that it is the first study to explore selective spatial responses for a naturalistic dynamic stimulus and task.

Earlier visual studies have applied similar approaches using EEG (Gonçalves et al., 2014; Lalor et al., 2006), but were limited to a discrete stimulus paradigm. We have shown previously that neural response to optical flow is enhanced during active play (Ki et al., 2020); however, we had not considered spatial heterogeneity of stimulus dynamics. We hope that the new approach and findings motivates future studies to explore complex relationships of neural activity with visual dynamics involving naturalistic stimuli. By further probing other spatial and temporal dimensions of the stimuli (beyond optical flow), or linking the regression to objects and features in the scene rather than locations, we may gain new insights into how neural signals convey visual processing during dynamic natural visual experiences.

## Methods

### Participant

In total, 42 healthy human subjects (18 females) aged 20 ± 1.56 participated in this experiment. All subjects provided written informed consent in accordance with procedures approved by the Institutional Review Board of the City University of New York.

### Video Stimulus

SuperTuxKart (an open-source game) is a Super Mario kart-like video game that emulates driving experience. All experimental trials were conducted on the default course and spanned three laps in “easy” mode. Game-related graphical interfaces were programmatically altered from the original game in order to minimize the visual elements and to simplify the task. The modified version of the game only involved the race car, track, passive objects on the road, and opponents.

### Experimental Procedure

In the original data collection, subjects played the game in 4 different task conditions, which is described in (Ki et al., 2020). Here, in the subsequent analysis, we only report trials from “active play” and “passive viewing” conditions. During active play, the subject’s used a keyboard to control the kart (using right hand): the left and right keys-controlled steering, while the up and down keys produced acceleration and braking. During passive viewing, subjects freely viewed a playback of previously recorded games. The recordings shown in passive viewing were distinct and not previously seen by the participants but were reused across participants (within each condition, all subjects viewed the same two stimuli). For each condition, subjects performed 2 trials. The ordering of all conditions was randomized and counterbalanced for all subjects. Prior to the recording of data, subjects were given an instruction on how to play the game, and it was followed by a practice trial.

### EEG Acquisition

The scalp electroencephalogram (EEG) was acquired with a 96-electrode cap (custom montage with dense coverage of the occipital region) housing active electrodes connected to a Brain Products ActiChamp system and Brain Products DC Amplifier (Brain Vision GmbH, Munich, Germany). The EEG was sampled at 500 Hz, digitized with 24 bits per sample.

### Video Capture Acquisition

The video game was played on a high-definition Monitor (Dell 24inch UltraSharp, 1920 by 1080 pixels, 60Hz) at a viewing distance of 60 cm in a dark and sound-dampened room with playback sound muted. The video game was rendered at 30 Hz. For each trial, the video frame sequence was captured with the Open Broadcaster Software (open-source), which records the screen display in real-time at the native resolution and frame rate.

### Eye Tracking Acquisition

Eye gaze (right eye) was measured using EyeLink 2000 (a video-based two-dimensional eye-tracking device, SR Research, Mississauga, Ontario, Canada). For every trial, the eye tracker was calibrated with a 5-point grid (center and four corners of the display). Subjects were asked to fix their gaze (in turn) at each of 5 locations where the dots are presented. This step was repeated until the error between all 5 eye-tracking positions and dot location were less than 2°. Of the 42 subjects, eye tracking data was collected from 24 subjects. The eye data was collected for every subject during the first cohort of the experiment (18 of 18 subjects) and part of the second cohort (6 of 24 subjects). Eye-tracking data was not used in the original study (Ki et al., 2020), thus the collection of the eye data was discontinued in the second cohort. Of the 24 subjects, 7 subjects missed eye-tracking in one of the conditions, thus they were excluded in the condition comparison analysis. In total, we had eye tracking data from 17 subjects (active play: 39 trials, passive viewing: 32 trials).

### EEG, Eye Tracking and Video Synchronization

To synchronize the video stimulus with the EEG we create flash events on the screen with a 30-by-30 pixel square placed (programmatically implemented) at the top right corner of the game, which flickered on (white) and off (black) at ~2Hz from the start to the end of each trial. A photodiode attached to this square electronically transmitted the on-screen flicker events to a separate event data acquisition hardware that relayed the signal via parallel port to a recording computer (which collected all incoming data: EEG, Eye Tracker, photodiode). To synchronize the data, we used Lab Streaming Layer software (Kothe, 2014), which created timestamps (in real-time) for all incoming data under a universal clock.

To obtain the event markers of the video capture, we isolated the flickering square patch for individual frames and computed the mean pixel intensity values of this region—the pixel intensity events marked on and off flash events on-screen. Using the event markers from the video and the auxiliary channels, we synchronized the video capture with EEG and Eye Tracking data.

### Optical Flow Extraction

The video capture of each kart race game was downsampled at a spatial resolution of 320-by-180 pixels and temporal resolution of 30 frames per second using Video Convert Factory (WonderFox Soft Inc). The video is grayscaled and further downsampled to a 24-by-32 patch. The optical flow was computed using MATLAB Computer Vision System Toolbox (Horn & Schunck, 1981) and for each patch, optical flow is normalized (z-score). This yields 368 (24 height x 32 width) time series features, *s*(*t*) for each trial.

### EEG Processing

EEG data was high-pass filtered at 1 Hz to remove slow drifts and down-sampled to 30Hz to match the frame rate. We employed the robust PCA technique (Candès et al., 2011) to remove gross artifacts from the data using the implementation of Lin et al. (Lin et al., 2013) with the default hyperparameter of *λ* = 0.5. This robust PCA method provides a low-rank approximation to the data and thereby removes sparse noise from the recordings. To reduce the contamination of eye movement from EEG, we linearly regressed out the activity of four virtual electrodes constructed via summation or subtraction of appropriately selected frontal electrodes (F9, Fp1, Fp2, F10). To further denoise the EEG, we rejected electrodes whose mean power exceeded the mean of all channel powers by four standard deviations. Within each channel, we also rejected time samples (and its adjacent samples) whose amplitude exceeded the mean sample amplitude by four standard deviations. These samples were replaced with zeros. We repeated the channel and sample rejection procedures over three iterations.

### Eye Tracking Processing

The raw data was filtered by EyeLink2000 using set criteria for blinks and fixations, which were based upon gaze position and the velocities of the individual points. The eye-tracking data was downsampled to 30Hz to match the frame rate of the video. The gaze velocity is computed by differentiating each gaze point, *v*(*t*) = *x*(*t* + 1) − *x*(*t*). For further outliers and jitter artifacts removal, we find and replace samples (with not a number marker) in which the gaze velocity magnitude (squared magnitude of the x and y gaze position derivative) was 3 standard deviations greater than the gaze velocity magnitude for a given trial. For computing the gaze position, we compute a histogram across 2 dimension space, which shows the number of times eye gaze was directed at a patch location for individual trials. For display, we normalize and apply gaussian smoothing over the 2d histogram image.

### Spatially-Resolved Stimulus-Response Correlation (SRC)

The main goal of this study is to assess the difference in neural response to visual dynamics across the visual space. Specifically, we measure the temporal correlation of continuous EEG to optical flow magnitude across the screen. At each patch location on the screen, optical flow, *s*(*t*) is linearly mapped to neural response, *r*(*t*) via temporal response function (Crosse et al., 2016; Lalor et al., 2006). This relationship is defined by simple linear convolution, 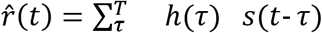, where *h*(*τ*) is the temporal coefficients at each time lag (*τ*) with temporal window length, *T* = 30 (equivalent to the video sampling rate). In this study, neural response can be represented by either individual EEG electrode *r_i_*(*t*), where *i* denotes a channel index (Figure 4), or ‘virtual electrodes’ 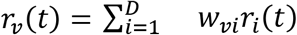, where, *r_v_*(*t*) represents a linear combination *D*-electrodes with spatial weights, *w_i_* across the scalp (Figure 3). The temporal response filter *h*(*τ*) is computed via ridge regression, separately for each (virtual or actual) electrode. Stimulus response correlation (SRC) is then the correlation between the estimated response 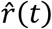 and the actual neural response *r*(*t*). For Figure 6, we summed this SRC across the three virtual components of Figure 3.

The virtual electrodes, i.e., spatial EEG filters, *w_vi_* shown in Figure 3 are previously computed in Ki et al 2020 using canonical correlation analysis (Dmochowski et al., 2018) using the global mean-optical flow without resolving by location. The virtual electrodes are represented by the forward model *a_j_*, which is the equivalent of a “spatial response function” (Haufe et al., 2014; Parra et al., 2005).

### Statistical significance of SRC

To test the significance of SRC, we compare the probability of SRC for all trials to a set of 100 phase-randomized data. We compute the probability of SRC at individual patches being greater than the SRC computed on the surrogate data. The patch locations that are below chance significance (p > .05) is greyed out.

### Statical comparison between active play and passive viewing

For each condition, we computed SRC for every trial (active play: 84 trials, passive viewing: 84 trials) across the 24 x 32 patch location (768 patch locations in total). To compare the statistical difference between the active play and passive viewing trials, we performed Wilcoxon rank-sum test and showed the patches of significant difference (p < .05) by drawing white pixels over the patch. To correct for multiple comparisons, we controlled the false discovery rate at 0.05 across the 768 patch regions.

For comparing average gaze position and saccade velocity, we account for the missing trials by counterbalancing the samples. Here, we averaged the trials for individual subjects, resulting in 17 samples for each condition. For this, we apply pairwise comparison (Wilcoxon signed-rank), and for gaze position, we correct for multiple comparisons across the patches.

## Acknowledgments

We thank Bart Krekelberg, Timothy Elmore, Jay Edelman, Simon Kelly for comments on earlier versions of this manuscript. This study was supported by a grant from the U.S. Army Research Laboratory (ARL/DSO W911NF-10-2-0022).

## Supplement

**Supplementary 1.**
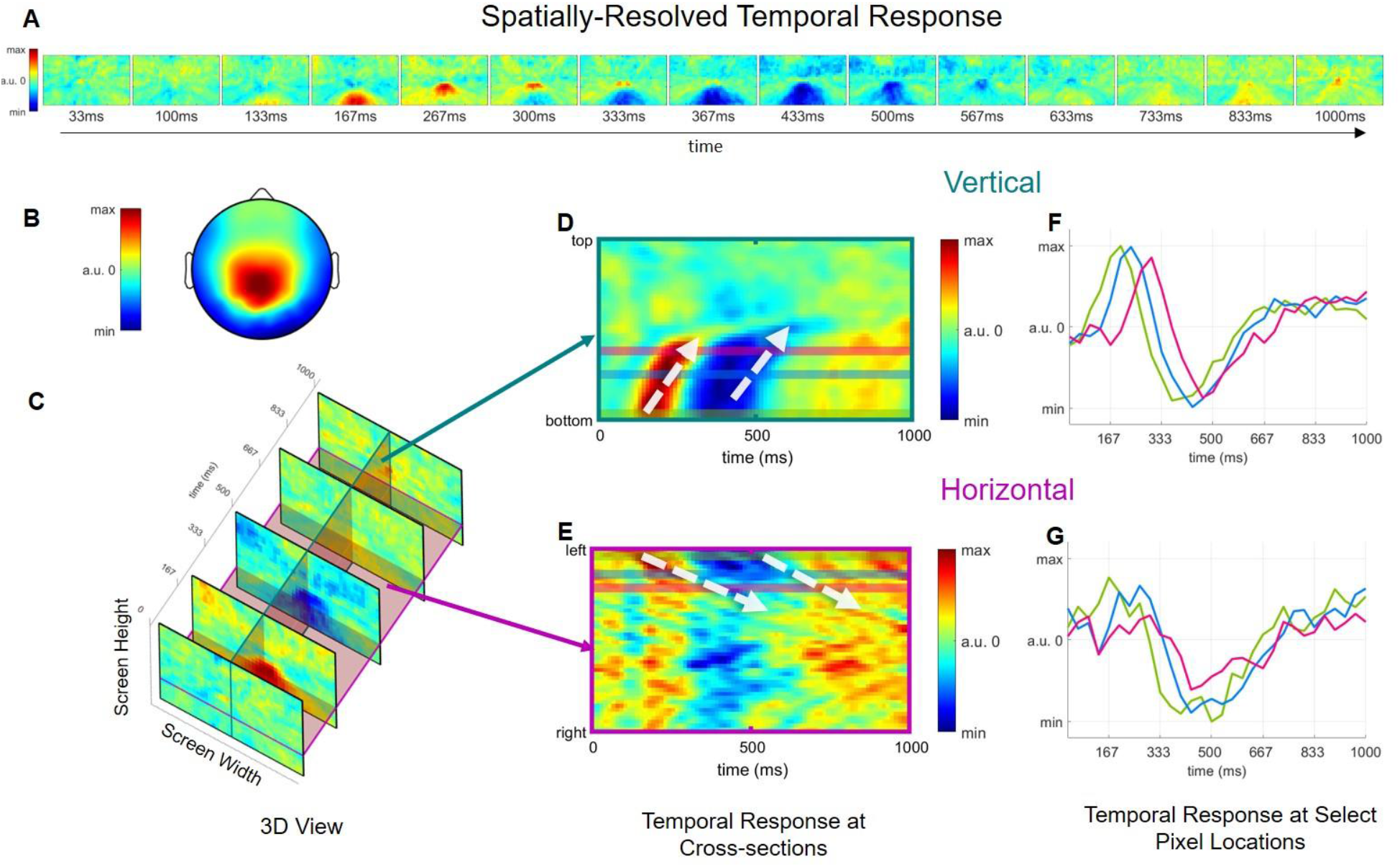
Spatially Resolved TRF of virtual electrode (component 1). Same as in figure 4 but for ‘virtual electrode’ component 1 (Figure 3).

**Supplementary Figure S2:**
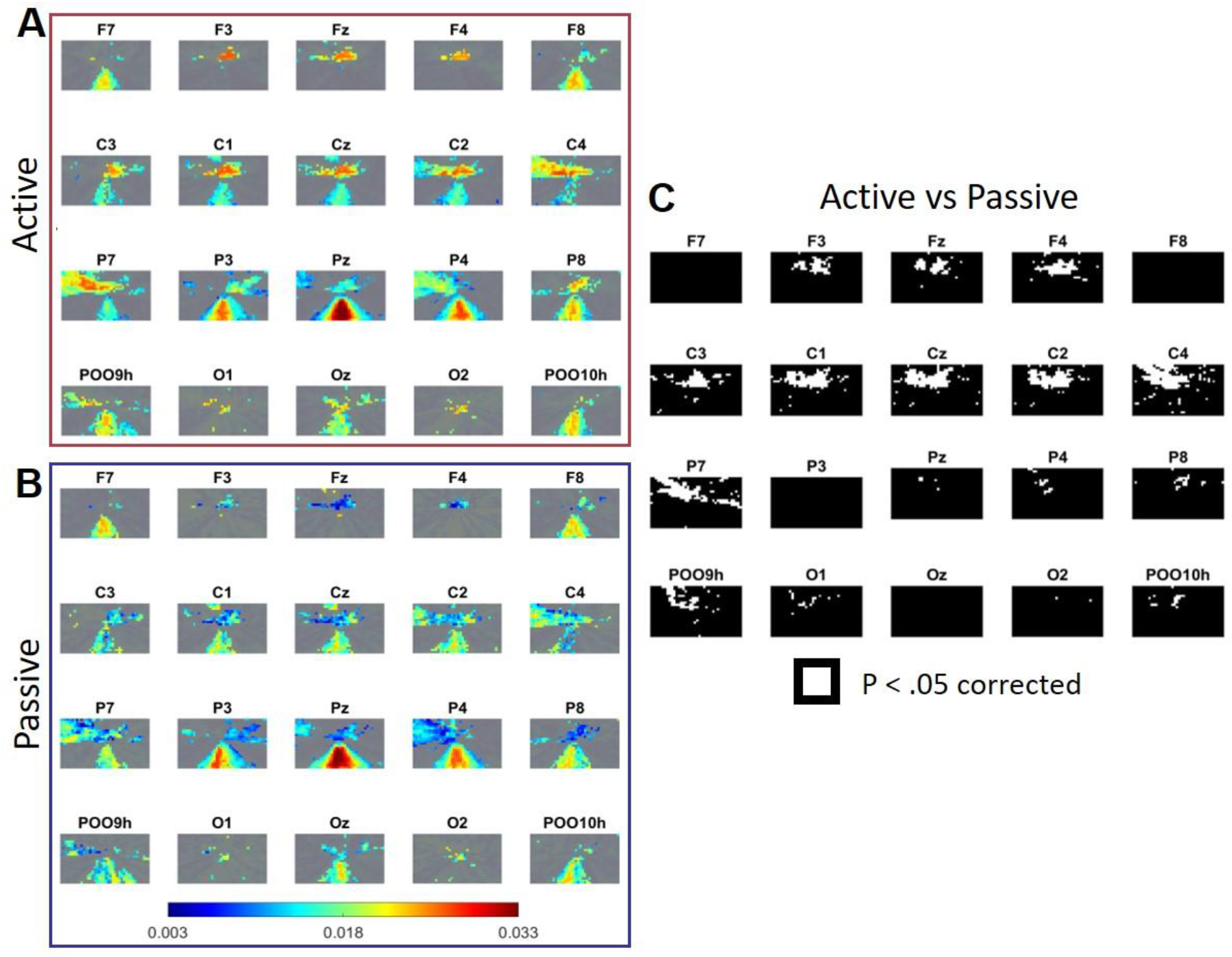
Same as figure 4, we computed spatially-resolved SRC for individual electrodes for each task condition: **A**. active play and **B**. passive viewing. **C.** Patches of significant differences in SRC between active and passive conditions for individual electrodes (P < .05, FDR corrected).

**Supplementary Figure S3:**
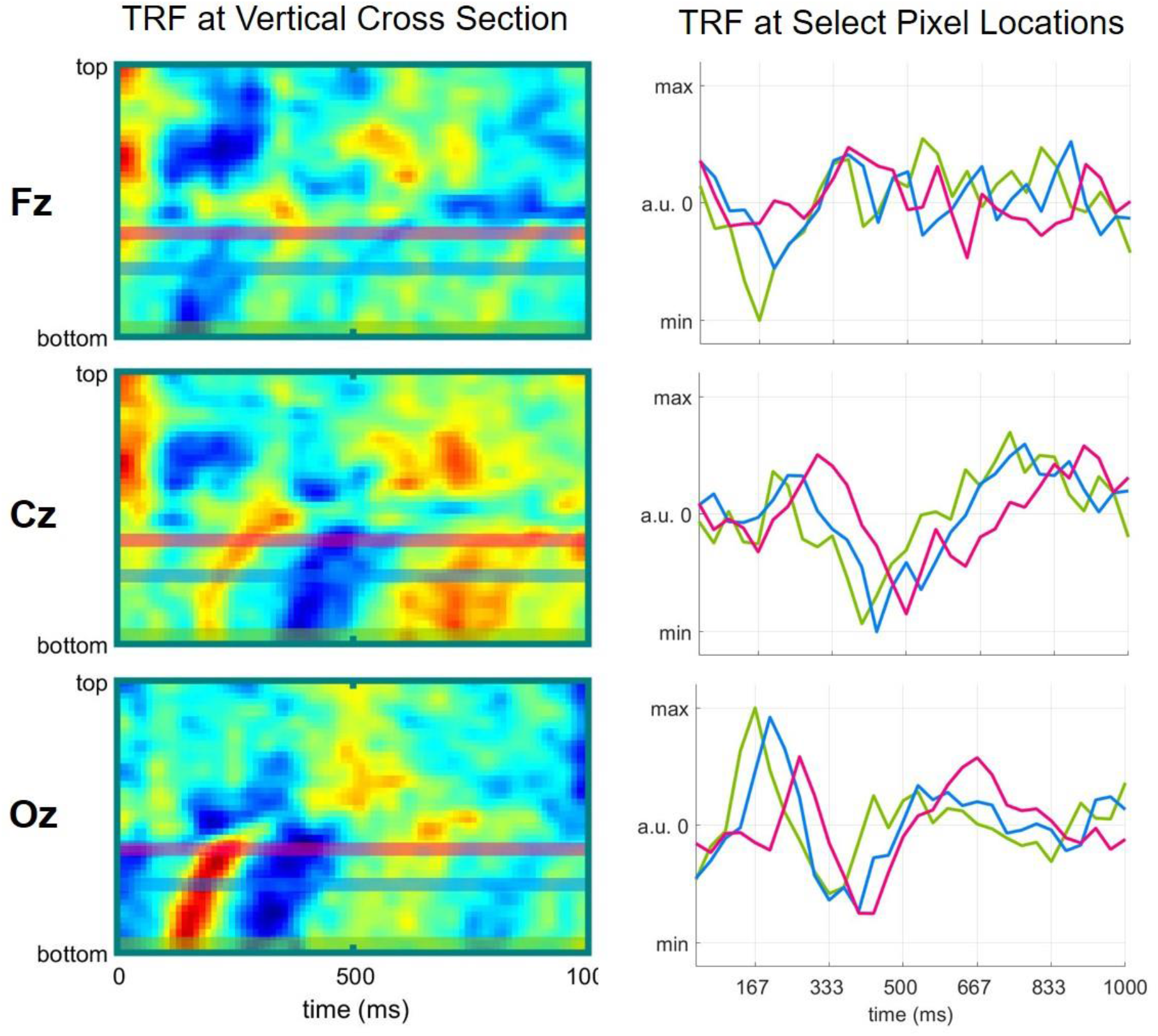
Temporal response for Fz, Cz and Oz electrodes across the vertical cross section.

**Supplementary Figure S4.**
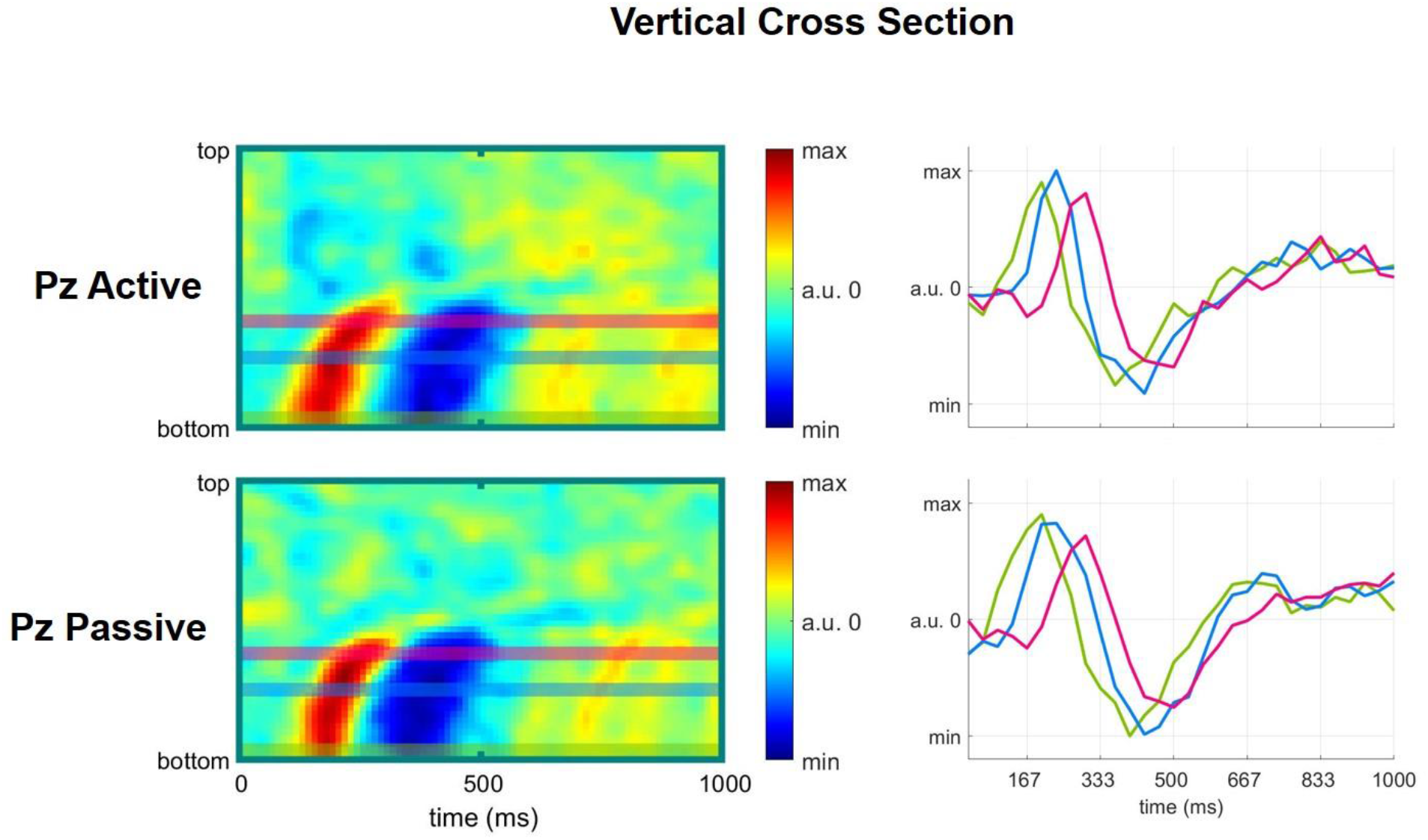
Comparison of spatially-resolved temporal response at a vertical cross section between active and passive conditions for Pz electrode.

## Notes

https://drive.google.com/file/d/1xt7OvM2xuVep_C-MEGd9cmqnQ3XHMxvO/view

### Competing Interest Statement

The authors have declared no competing interest.

